# Individual Host Variation in Single-cell Responses to Enteroaggregative *Escherichia coli* Infection Modeled with Human Colon Organoids

**DOI:** 10.1101/2025.10.24.684350

**Authors:** Ling Qiu, Tajhal D. Patel, Hannah E. Jones, Cristian Coarfa, Anthony W. Maresso

## Abstract

Enteroaggregative *Escherichia coli* (EAEC) is a leading cause of diarrhea disease worldwide. Interestingly, EAEC disease outcomes vary dramatically among infected individuals, ranging from asymptomatic colonization to acute and chronic diarrhea. While previous research has focused on EAEC virulence factors, emerging evidence suggests that host genetic and cellular background may be equally important in determining disease susceptibility. To understand host-driven disease heterogeneity, we performed single-cell RNA sequencing on human colon organoids (HCOs) derived from three healthy adult donors following EAEC 042 infection. Surprisingly, despite comparable EAEC adherence across donors, we observed striking donor-specific responses to infection at multiple levels. A total of seven distinct colonic epithelial cell clusters were identified, with striking donor-specific baseline compositions. Following infection, the HCO donors exhibited three distinct response phenotypes evident in both cell type remodeling and gene regulation: hyperresponsive, intermediate, and non-responsive. We further observed extensive donor-specific cell-cell communication networks at baseline and after EAEC infection. This study suggests that the heterogeneity in host epithelial response to EAEC infection extends to the single-cell level. These findings establish a paradigm for understanding infectious disease susceptibility through the human organoid model system.

**IMPORTANCE:** Understanding why similar pathogen exposures produce vastly different clinical outcomes is critical for combating infectious diseases. By conducting single-cell RNA sequencing on human colon organoids derived from three different donors, we show that each donor possesses unique intrinsic cellular compositions and distinct transcriptomic responses to EAEC infection. These findings challenge the traditional pathogen-focused paradigm and highlight the critical role of host factors in infectious disease outcomes. This study also validates human organoids as a powerful platform for studying host heterogeneity, which may help predict disease susceptibility and optimize treatment for diverse patient populations in the future.

## INTRODUCTION

Historically, research on bacterial pathogenesis has largely focused on the pathogens. In particular, studies in the field have emphasized bacterial virulence factors involved in nutrient acquisition, colonization, toxin production, and immunomodulation^1^. While such research has led to successful vaccines and anti-virulence therapeutic strategies, it often overlooks an equally critical component of disease: the host. Host factors – including age, sex, genetics, microbiome, immune status, and comorbidities – play a central role in determining susceptibility to bacterial infection and disease severity^2^. Leveraging host biology offers great potential for tackling infections and addressing the growing threat of antibiotic resistance. However, our understanding of host contributions to pathogenesis remains severely limited.

Enteroaggregative *Escherichia coli* (EAEC) is a group of enteric bacteria that cause persistent diarrhea in children^3^, travelers^4^, and immunocompromised patients^5^. Pediatric EAEC infections are particularly problematic due to its association with malnourishment and impaired growth^6^, even in the absence of overt diarrhea symptoms^7^. Despite its global impact and serious sequalae, treatment options for EAEC remain limited to ineffective antibiotics, largely due to widespread multidrug resistance observed in over 70% of clinical isolates^8^. To combat the emergence of highly virulent strains, a deeper understanding of the molecular mechanisms underlying EAEC pathogenesis is essential. Major EAEC virulence factors include aggregative adherence fimbriae (AAF), which mediates adhesion to and colonization of the intestinal mucosa; the protein involved in colonization (Pic), which degrades mucin; and the plasmid-encoded toxin (Pet), which induces cytotoxicity^9–10^. Nevertheless, clinical manifestations of EAEC infection range widely – from asymptomatic carriage to acute and chronic diarrhea – a variability not fully explained by EAEC virulence factors or bacterial load^11–12^. Furthermore, human volunteer studies have shown that the ability of EAEC to colonize and cause diarrhea varies depending on the host^13^. These findings further highlight the importance of the host in determining infection outcomes and underscore the urgent need to better understand host biology in the context of EAEC pathogenesis.

Using EAEC as a pathogen to model host differences, we aim to investigate host-specific variation in infection response using human colon organoids (HCOs). HCOs are non-transformed, tissue stem cell-derived models that, upon *in vitro* differentiation, closely recapitulate the cellular composition and functional characteristics of the mature human colon epithelium^14^. Crucially, HCOs retain the genetic diversity of their donors, offering a unique opportunity to examine how host genetic background influences infection outcomes. Previous research from our group^15–16^ and others^17^ has established HCOs as a robust model for studying EAEC infection, revealing both segment- and host-specific patterns of colonization. However, while bacterial binding and colonization patterns have been explored, differences in the host response to EAEC infection across donors remain largely unexamined. Recent studies have leveraged single-cell RNA sequencing (scRNA-seq) to dissect cell type–specific responses to pathogens within the intestinal epithelium^18–20^. However, most scRNA-seq studies involving human organoids treat donor variation as a source of biological noise, averaging out potentially meaningful differences. By harnessing the inherent genetic diversity of HCO donors, we investigated donor-specific infection responses at single-cell resolution. This approach helps uncover epithelial cell subtypes and pathways that confer protection against EAEC, advancing our understanding of host determinants in bacterial pathogenesis. Here, we describe substantial host-specific response at the single-cell level despite uniform infection conditions and pathogen strain.

## RESULTS

Building off previous results^15–16^ that the response to EAEC infection is driven largely by differences in the host, we sought to determine if such observations extend to individual cells within patient-derived organoid lines. To test this idea, we selected three HCO lines from our biobank and performed EAEC infection followed by single cell RNA sequencing. Organoids from the colon were used because EAEC shows a strong large bowel disease presentation^21^. For clarity, we refer to these lines as X, Y, and Z and report the donor characteristics in **Supplementary Table 1**. The experimental workflow is illustrated in **Figure 1A**. In brief, colonoid monolayers were differentiated for 4 days, then either mock-infected or infected with EAEC strain 042 for 1.5 hours. Following infection, the monolayers were then dissociated into single cell suspensions and subjected to single cell RNA-sequencing using the 10X Genomics platform. To ensure comparable levels of infection across lines, we quantified bacterial burden by determining colony-forming units (CFUs) adhering to the monolayers after the washing step. As shown in **Figure 1B**, there were no significant differences in bacterial adhesion among the three donor lines, indicating that any variation in host response is likely due to intrinsic differences in host genotype or phenotype, rather than differences in bacterial load.

**Figure 1.**
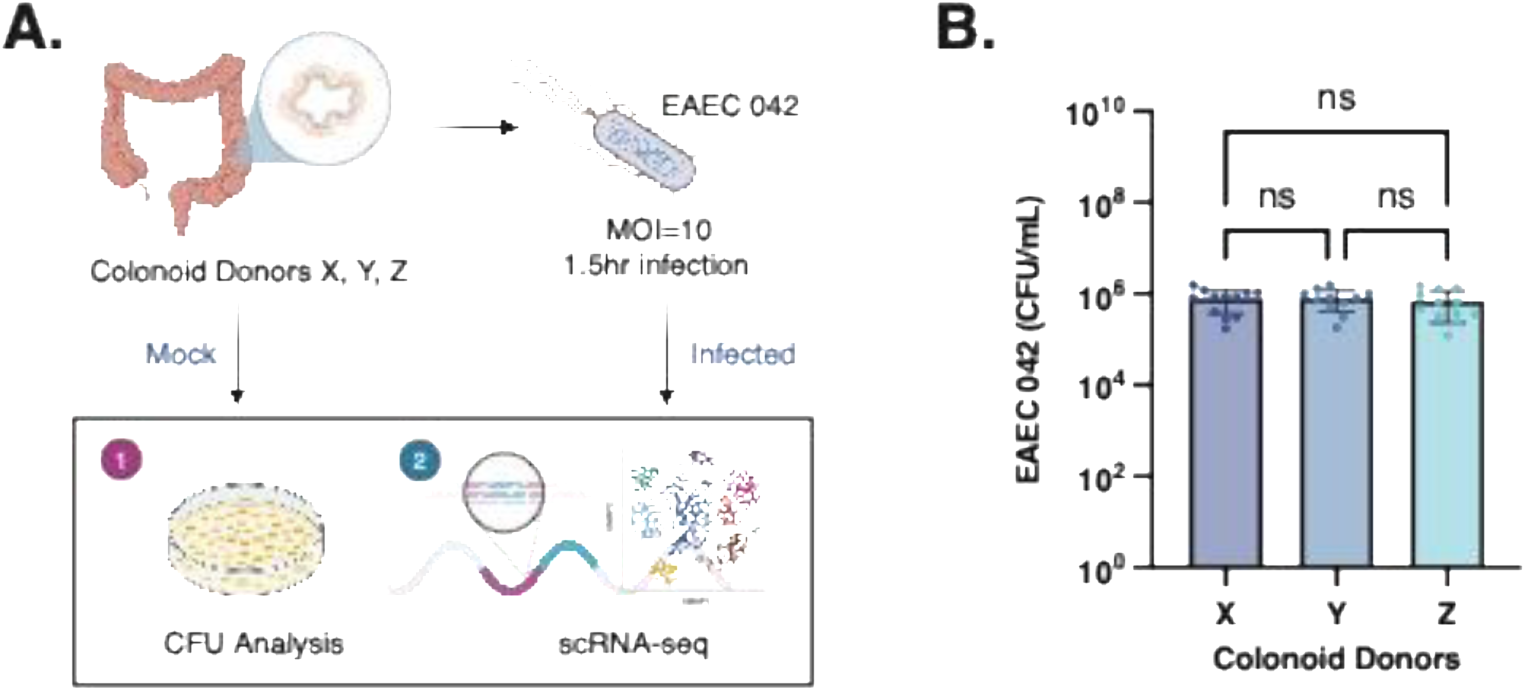
Colonoid scRNA-seq Experiment Design. **(A)** Schematic of scRNA-seq and bacterial quantification on EAEC-infected colonoids. Briefly, colon organoids derived from intestinal crypts of 3 healthy donors were plated as monolayers on a 96-well plate and differentiated for 4 days. Colonoid monolayers were either mock infected with fresh differentiation media or with log phase EAEC 042 at MOI=10 for 1.5 hours at 37°C in a humidified incubator. After infection, colonoid monolayers were washed to remove unbound bacteria, collected and separated into single cell suspensions, and then sequenced via the 10x platform. Additional colonoids were collected and plated on LB agar to quantify bacterial loads. **(B)** Bar plot depicting EAEC 042 load (CFU/mL) adhered on colonoid monolayers after 1.5 hours of infection.

### Cell Type Annotation and Donor Intrinsic Differences

A critical first step before evaluating EAEC-induced transcriptional changes is to accurately annotate the cellular composition of the population. After QC filtering, a total of 39,351 cells remained for downstream data analysis. Filtered scRNA-seq data including all mock and infected cells, was first processed using Scanpy and Seurat. Clustering with the Leiden algorithm^22^ revealed 7 distinct cell populations, visualized by Uniform Manifold Approximation and Projection (UMAP) (**Figure 2A**). We next annotated these clusters as 7 different colonic epithelial cell subtypes, including colonocytes, goblet cells, and precursor cells (**Figure 2B**). To define the developmental relationships among the precursor populations, we applied Slingshot pseudotime analysis (**Supplementary Figure 1A-B**). This analysis suggested a differentiation trajectory originating from Precursor Cells 1, the least differentiated population, progressing through Precursor Cells 2 and Precursor Cells 3, before diverging into 4 differentiated populations: *BEST2⁺* Goblet Cells, *CA2⁺* Colonocytes, *CA1⁺* Colonocytes, and *CEACAM7⁺* Colonocytes. The naming of Precursor Cells 1, 2, and 3 reflects their inferred order along this developmental trajectory. Each precursor cluster expressed distinct combinations of proliferative and epithelial lineage markers (**Figure 2B**). Precursor Cells 1 were enriched for the progenitor marker *POLR2J3* and early enterocyte markers *RSRP1*, *VMP1*, and *KLF6*. Precursor Cells 2 expressed intestinal stem cell markers *RPL12* and *PCNA*, alongside the colonocyte marker *CA2*. Interestingly, Precursor Cells 3 co-expressed proliferative markers (*RPL12*, *GDF15*, *PCNA*), goblet cell markers (*MUC4*, *ZG16B*, *SPINK1*, *RAMP1*), tuft cell markers (*KRT7*, *MYO1B, TRIM29*, *SLC6A14*). This population also showed high expression of chemokine-encoding genes including *CXCL1*, *CXCL8*, *CXCL3*, and *CXCL16*. All three colonocyte populations expressed canonical enterocyte markers *FABP2* and *SLC26A2* but were distinguished by cluster-specific signatures. *CA1⁺* Colonocytes expressed *CA1*, *PDZK1*, *FAM3B*, *MTTP*, and *NMU*; *CA2⁺* Colonocytes expressed *CA2*, *LAMA1*, *LINC00668*, *GABRA2*, and *RPS4Y1*; and *CEACAM7⁺* Colonocytes expressed *CEACAM7*, *KCNQ1OT1*, *BTNL8*, *SLC30A10*, and *PLOD2*.

**Figure 2.**
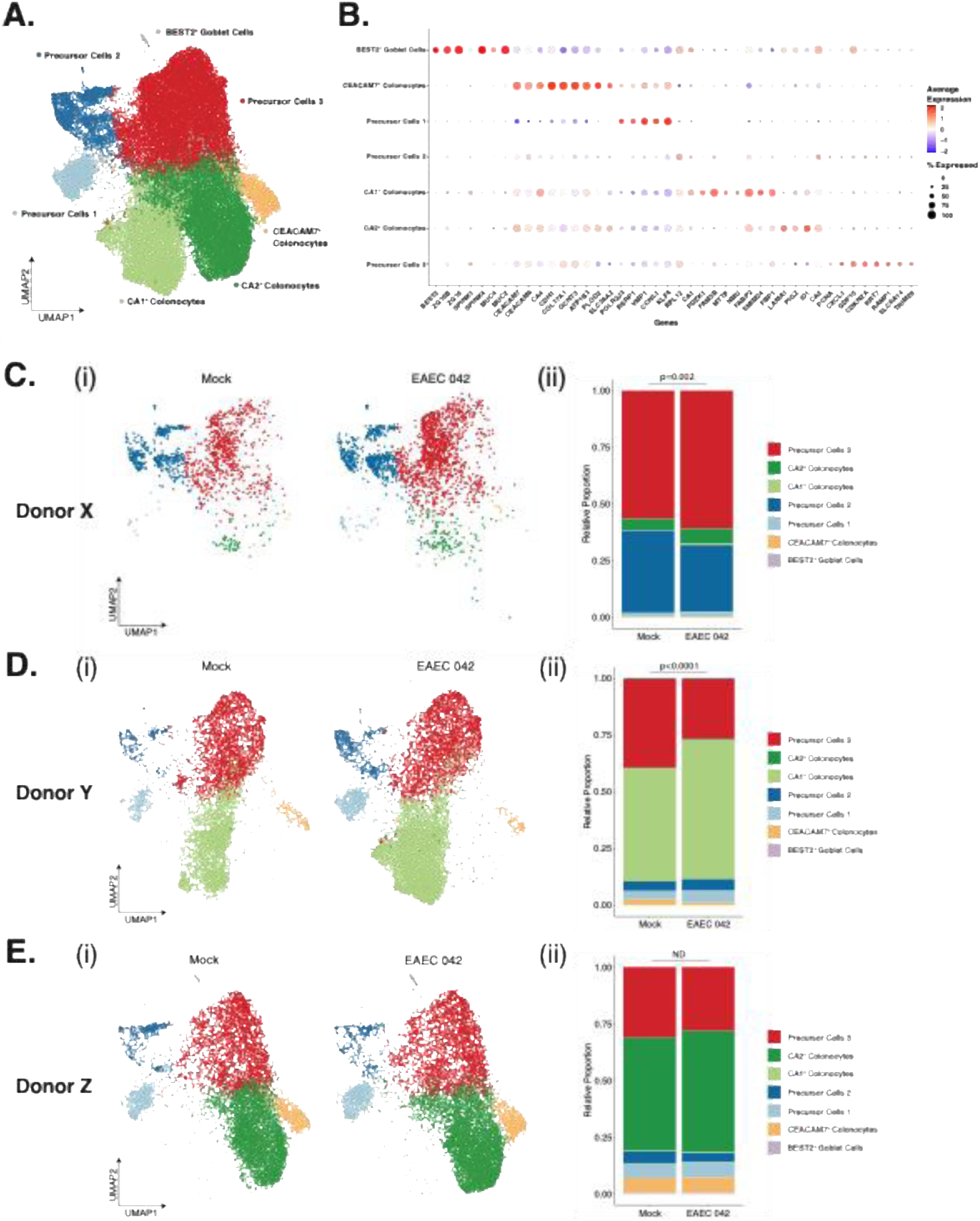
Donor-specific cell type distribution and changes induced by EAEC infection. **(A)** Uniform Manifold Approximation and Projection (UMAP) showing 7 cell clusters at Leiden resolution 0.4. **(B)** Dot plot showing relative gene expression of cell cluster markers. **(C)** Cell cluster distribution of Donor X under mock- or EAEC-infection conditions, visualized using UMAP **(i)**, with the relative proportion of each cluster presented in the accompanying bar plot **(ii)**. **(D)** Same as **(C)** but for Donor Y. **(E)** Same as **(C)** but for Donor Z. Change in cell composition between Mock and EAEC is assessed using a Chi-Square test.

Next, we stratified the 7 colonic epithelial cell clusters by donor and observed distinct cell type distributions across the three colonoid lines (**Supplementary Figure 1C-D**). Donor X exhibited a predominance of undifferentiated populations, with over 90% of cells classified as either Precursor Cells 2 or Precursor Cells 3. In contrast, differentiated cell types were enriched in Donors Y and Z, with *CA1⁺* Colonocytes being the major population in Donor Y, and *CA2⁺*Colonocytes dominating in Donor Z. Notably, *CA1⁺* Colonocytes were largely unique to Donor Y, while *CA2⁺* Colonocytes and *BEST2⁺* Goblet Cells were almost exclusively observed in Donor Z, highlighting striking donor-specific intrinsic differences in epithelial cell composition.

### Infection-driven Differences in Cell Clusters

Evaluating changes in cell type distribution induced by EAEC 042 infection for each donor, we were surprised to find significant host-specific changes (**Figure 2C-E**, **Supplementary Figure 1E-F**). Among the three donors, Donor Z appeared relatively resistant to EAEC colonization (**Figure 2E**), exhibiting no significant changes in cell type composition following infection – classifying this donor as a non-responder to EAEC infection. In contrast, both Donors X and Y displayed significant alterations in cluster distribution after bacterial exposure (**Figure 2C-D**), therefore classifying as responders to EAEC infection. In Donor X (**Figure 2C**), infection was associated with an increase in Precursor Cells 3, a population enriched in immune-activating transcripts. Donor Y (**Figure 2D**), on the other hand, showed an enrichment of CA1⁺ Colonocytes and a depletion of Precursor Cells 3, potentially reflecting enhanced epithelial function such as pH regulation by mature colonocytes. While the mechanistic basis for these shifts remains to be elucidated, our findings indicate that organoid donor identity not only influences susceptibility to EAEC adherence^15–16^ – a hallmark of EAEC pathogenesis – but also drives distinct transcriptional and cellular remodeling during infection at the single cell level.

### Infection-driven Differences in Gene Regulation

Since each donor displayed a “personalized” cell distribution upon infection, we next asked whether the cell populations within each donor also differentially regulate gene expression in response to EAEC infection. To address this, we performed a differentially expressed genes (DEG) analysis (Mann-Whitney-Wilcoxon, FDR<0.1, fold change exceeding 1.25x). As an initial step, we confirmed that the observed differences could not be attributed to baseline gene expression variation among donors (**Supplementary Figure 2**). Overall, a greater number of genes were downregulated than upregulated (**Figure 3A**). The majority of DEGs arose from *CA1^+^* Colonocytes in Donor Y, followed by *CA2^+^* Colonocytes in Donor X, Precursor Cells 3 in Donor Y, and *CEACAM7^+^* Colonocytes in Donor Z. Interestingly, While Donor X displayed mostly upregulated DEGs, Donors Y and Z exhibited a predominance of downregulated DEGs in response to infection.

**Figure 3.**
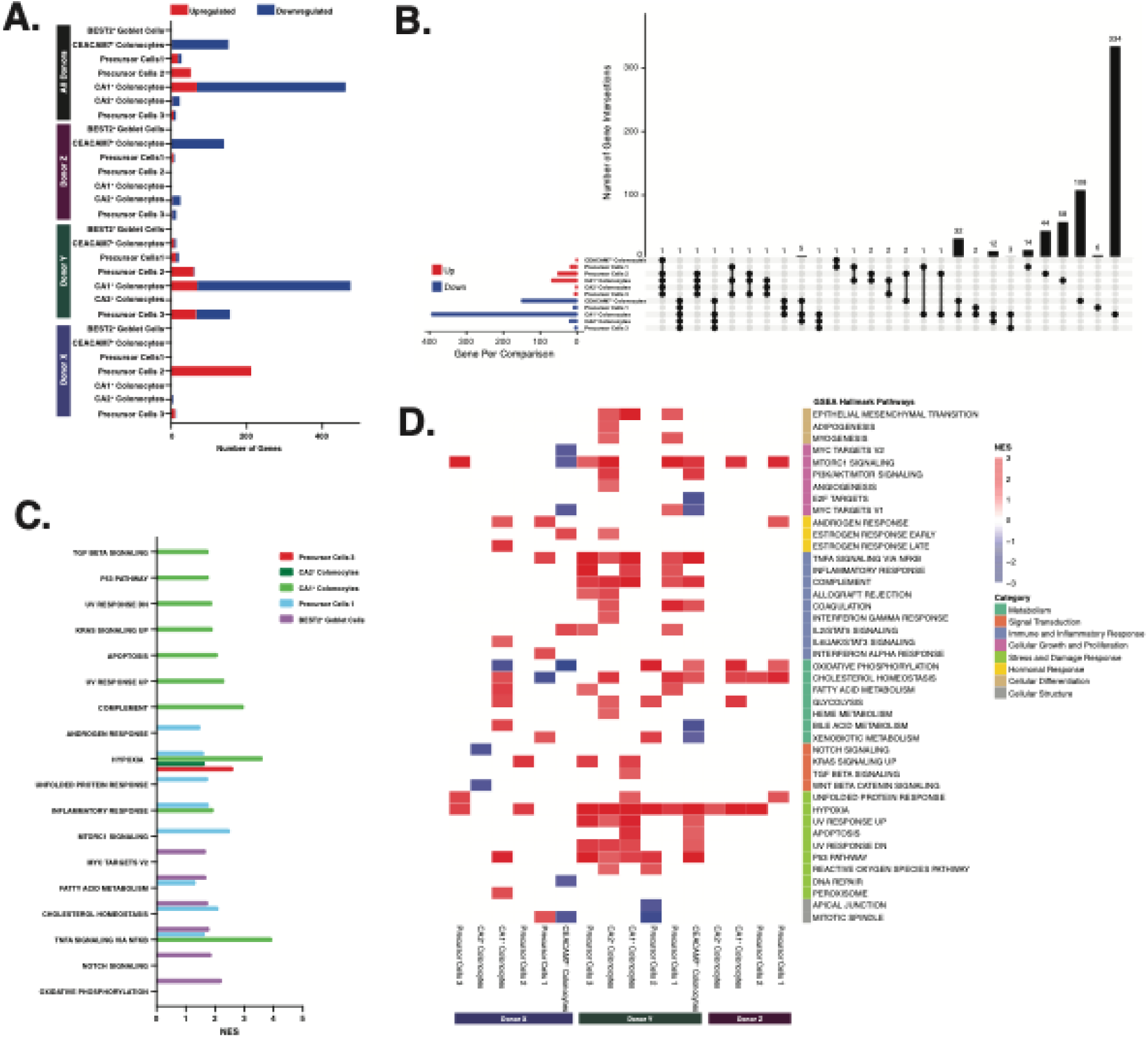
Donor-specific gene and pathway regulation induced by EAEC infection. **(A)** Stacked bar diagram showing numbers of differentially expressed genes (DEGs) between EAEC-infected and mock-infected colonoids (FDR<0.1, fold change exceeding 1.25x). **(B)** UpSet plot showing overlapping DEGs across different cell clusters using pooled donors. **(C)** Bar plot showing infection-enriched hallmark pathways for different cell clusters, using Gene Set Enrichment Analysis (GSEA). Normalized Enrichment Scores (NES) are presented. **(D)** Heatmap showing infection-enriched hallmark pathways for each cell cluster within each donor. NES are shown for enriched pathways.

We next examined overlap among DEG sets across clusters computed by pooling all the donors (**Figure 3B**) and found that most DEGs were cluster-specific, suggesting that each population plays a distinct role in responding to infection. Nonetheless, several patterns of convergence emerged: (1) the three colonocyte clusters shared five commonly downregulated genes, (2) *CA1^+^* and CEACAM7^+^ Colonocytes shared 32 downregulated DEGs, and (3) *CA2^+^* and *CA1^+^*Colonocytes shared 12 downregulated DEGs. These findings highlight that although donors harbor different proportions of colonocyte subtypes, these cells share conserved transcriptional programs to combat infection.

To further investigate molecular pathways altered by EAEC infection, we performed GSEA hallmark pathway analysis for the infection response. Overall, we observed a general upregulation of hallmark pathways (**Figure 3C**), primarily driven by *CA1^+^* Colonocytes, Precursor Cells 1, and *BEST2^+^*Goblet Cells. Notably, most upregulated pathways were associated with immune and inflammatory responses, as well as stress and damage responses. For instance, hypoxia pathways were elevated in four clusters, and TNF*α* signaling via NF-kb pathways were enhanced in three clusters.

When examining pathway regulation by donor (**Figure 3D**), we found striking variability in their responses to EAEC infection. Donor Y exhibited the strongest response, with widespread upregulation in pathways related to cellular differentiation, immune and inflammatory signaling, metabolism, and stress responses. By contrast, Donor Z showed the weakest response, with no detectable changes in cellular differentiation, immune response, signal transduction, or structural pathways; this is consistent with the previous observation in the cell type analysis. Donor X displayed an intermediate level of responsiveness.

Interestingly, cell-type-specific responses varied by donor. For example, *CA2^+^* Colonocytes exhibited consistent upregulation of hallmark pathways in Donors Y and Z but displayed downregulation of signal transduction pathways in Donor X, specifically Notch and Wnt/*β*-catenin signaling. Similarly, *CEACAM7^+^* Colonocytes in Donor X showed broad pathway downregulation, including in MYC signaling, mTORC1 signaling, oxidative phosphorylation, and DNA repair. However, this cell cluster showed mainly pathway upregulation in Donor Y and no detectable changes in Donor Z. Together, these findings underscore that EAEC elicits highly donor-specific gene regulation events, with variability evident not only across individuals but also at the level of individual cell populations.

### Donor-specific cell-cell communication in response to infection

CellChat is a computational framework that enables the characterization of intercellular communication dynamics by analyzing ligand–receptor pairings in complex single-cell datasets^23^. Using this approach, we identified >1,500 significantly altered signaling interactions following EAEC infection (**Figure 4A**). Among the three donors, Donor Z exhibited the largest number of both upregulated and downregulated interactions. In contrast, Donor Y displayed the fewest increased signaling events, while Donor X exhibited the fewest decreased events. Strikingly, only a limited number of interactions were commonly altered across all donors, with 20 shared upregulated and 2 shared downregulated signaling interactions. These results suggest that individual donors exhibit not only distinct cell-specific responses to infection, but also different functional consequences of these responses.

**Figure 4.**
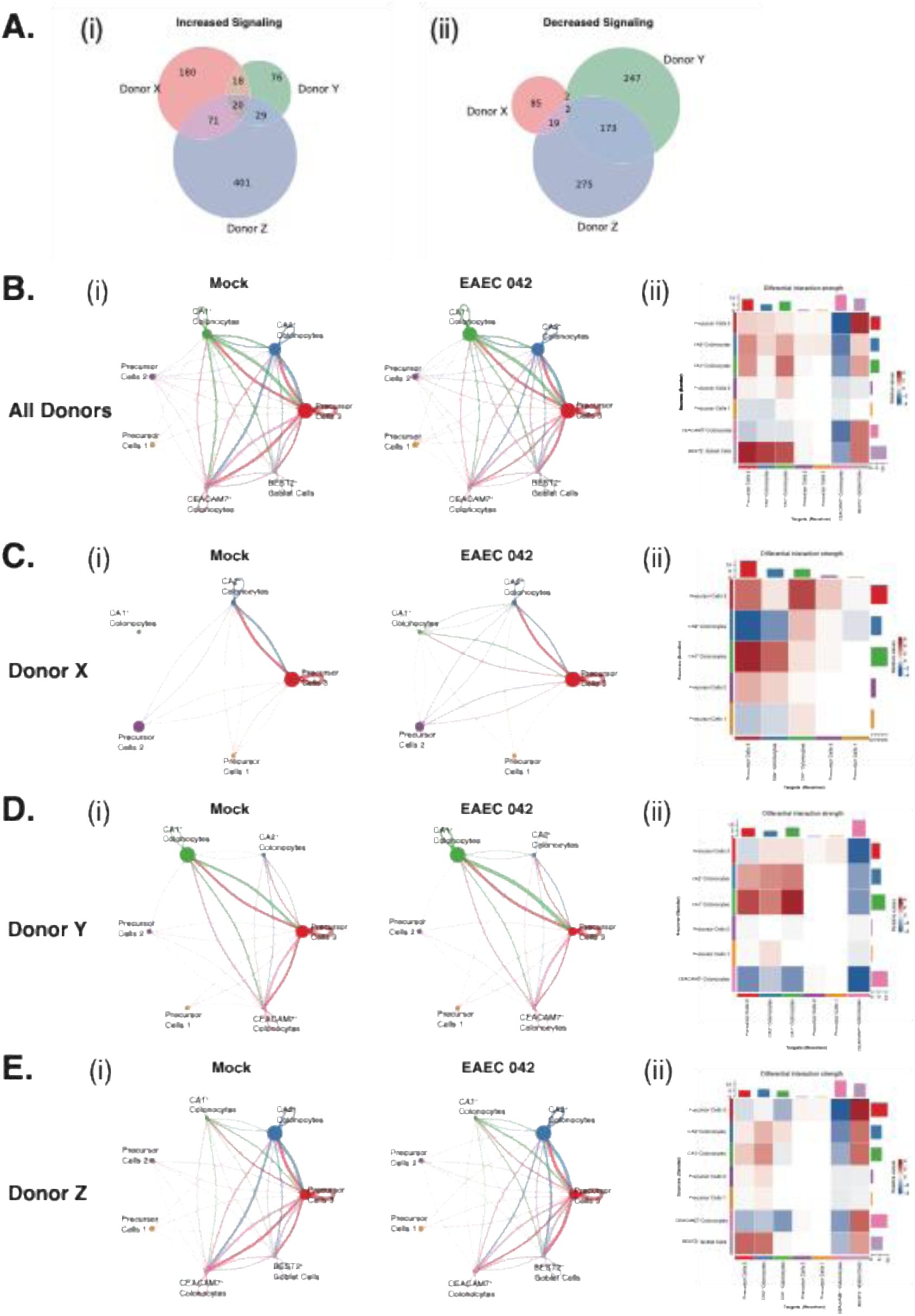
CellChat ligand-receptor analysis. **(A)** Venn diagrams showing the numbers of **(i)** increased and **(ii)** decreased signaling interactions caused by EAEC infection in each donor. **(B) (i)** Circle plots showing interactions between clusters with or without EAEC infection for all cells. **(ii)** Heatmap summarizes changes in cell-cell interactions for each pair of source and target clusters. **(C)** Same as **(B)** but for Donor X. **(D)** Same as **(B)** but for Donor Y. **(E)** Same as **(B)** but for Donor Z.

When pooling all donors, circular interaction plots revealed that in both mock- and EAEC-infected states, the majority of intercellular communications were mediated by Precursor Cells 3, *CA2*^+^ Colonocytes, and *CA1*^+^ Colonocytes **(Figure 4B-i**). Among these, Precursor Cells 3 were the most active signaling population, engaging in bidirectional communication with goblet cells and all mature colonocyte subtypes. In the uninfected state, Precursor Cells 1 were unique in exclusively initiating signaling without receiving input from other clusters, and both Precursor Cells 1 and 2 lacked intra-cluster communication. Upon infection, the most prominent changes were observed in crosstalk between Precursor Cells 3 and *BEST2^+^* Goblet Cells (**Figure 4B-ii**). Goblet cells also displayed enhanced intra-cluster signaling and increased communication with both *CA1^+^* and *CA2^+^* Colonocytes. Conversely, *CEACAM7*+ Colonocytes were the major targets of downregulated communication, particularly reducing interactions initiated by Precursor Cells 3. Notably, despite their overall reduction in signaling as targets, *CEACAM7*+ Colonocytes increased communication into *BEST2*^+^ Goblet Cells after infection. Precursor Cells 1 also began receiving signaling from Precursor Cells 3 and *CA2*^+^ Colonocytes upon infection, although overall, Precursor Cells 1 and 2 remained relatively stable in their communication landscape. Collectively, these findings underscore the central role of Precursor Cells 3 and *BEST2*^+^ Goblet Cells in coordinating host responses to EAEC infection.

Having established the overall landscape of cell-cell communication across all samples, we next examined how these signaling dynamics varied among individual donors (**Figure 4C-E**). Donor X exhibited the simplest network of signaling events (**Figure 4C**). In this donor, the strongest interactions occurred between Precursor Cells 3 and *CA2*^+^ Colonocytes, regardless of infection status. Under mock conditions, *CA1*^+^ Colonocytes did not participate in any signaling. However, following infection, *CA1*^+^ Colonocytes received signaling input from all clusters – most prominently from Precursor Cells 3 – and initiated communication with all clusters except Precursor Cells 1, including strong intra-cluster signaling. Precursor Cells 3 and *CA2*^+^ Colonocytes showed reduced signaling to Precursor Cells 1, leaving the latter with only outward communication. Interestingly, although *CEACAM7^+^*Colonocytes were present in Donor X (**Figure 2C**), they were not engaged in any cell-cell signaling activities.

In Donor Y (**Figure 4D**), the most robust interactions were observed between Precursor Cells 3 and *CA1*^+^ Colonocytes, with the stress from EAEC infection further amplifying this communication. Consistent with global analyses, Donor Y exhibited more downregulated than upregulated signaling events after infection (**Figure 4A**), the majority of which mediated by *CEACAM7^+^* Colonocytes.

Donor Z displayed the most complex communication network and the highest number of altered signaling interactions (**Figure 4A&4E**). In this donor, the strongest baseline interactions occurred among Precursor Cells 3, *CA2*^+^ Colonocytes, *CEACAM7^+^* Colonocytes, and *BEST2*^+^ Goblet Cells. EAEC infection resulted in marked increases in bidirectional signaling between Precursor Cells 3 and *BEST2*^+^ Goblet Cells, alongside pronounced bidirectional decreases between Precursor Cells 3 and *CEACAM7^+^*Colonocytes. Similar to Donor Y, *CEACAM7^+^* Colonocytes mediated the majority of downregulated interactions, yet their communication into *BEST2*^+^ Goblet Cells increased post-infection.

Taken together, these results demonstrate that while Precursor Cells 3 consistently serve as a central signaling hub across donors, each donor possesses a unique baseline communication landscape that is differentially remodeled by EAEC infection. These donor-specific communication profiles highlight the remarkable heterogeneity of host responses at the intercellular level within patient-derived colonoids.

### Donor-specific ligand-receptor response in CEACAM and GDF pathways

With the total cell-cell interactions showing striking donor variability both at baseline and in infection-induced changes, we next sought to investigate specific CellChat pathway interactions involved in EAEC infection. Carcinoembryonic antigen-related cell adhesion molecules (CEACAMs) are immunoglobulin-related glycoproteins that regulate key cellular processes including cell adhesion, differentiation, proliferation, and survival^24^. Altered CEACAM expression has been implicated in gastrointestinal diseases such as inflammatory bowel disease and colon cancer. Given the importance of CEACAM molecules in maintaining intestinal epithelial homeostasis, we investigated CEACAM-mediated communication across the seven colon epithelial cell subtypes. In this pathway, CEACAM1 functions as the ligand and CEACAM5 as the receptor. While both are broadly expressed in colon tissue, CEACAM1 is enriched in absorptive colonocytes, whereas CEACAM5 is more highly expressed in mucus-secreting colonocytes^25^.

In Donor X (**Figure 5A**), under mock-infected conditions, CEACAM signaling was primarily initiated by *CA2^+^* colonocytes and received by the same cell type as well as Precursor Cells 2 and 3. Following EAEC 042 infection, communication from *CA2^+^* colonocytes increased towards all these populations, and additional bidirectional signaling emerged between *CA2^+^* and *CA1^+^* colonocytes. Meanwhile, self-signaling within Precursor Cells 3 disappeared, and was replaced by input from *CA1^+^* colonocytes. In Donor Y (**Figure 5B**), *CEACAM7^+^* and *CA1^+^* colonocytes dominated signaling under both mock and infected conditions. *CEACAM7^+^* Colonocytes communicated with all cell types except Precursor Cells 1, while *CA1^+^* Colonocytes signaled broadly except to Precursor Cells 1 and 2. Unlike Donor X, where signaling largely increased post-infection, Donor Y exhibited a mixed pattern of gains and losses. Specifically, *CA2^+^* colonocytes strengthened communication with *CEACAM7^+^* colonocytes and Precursor Cells 3, while also establishing new interactions with *CA1^+^* colonocytes and their own population. By contrast, signaling from Precursor Cells 3 to *CEACAM7^+^* colonocytes was lost. In Donor Z (**Figure 5C**), *CEACAM7^+^* colonocytes were the primary signaling cluster, targeting nearly all clusters except Precursor Cells 1. *CA2^+^* colonocytes were also highly active, signaling to their own cells, Precursor Cells 3, *CEACAM7^+^*colonocytes, and *BEST2*^+^ goblet cells. EAEC infection further amplified these communications in Donor Z, while interactions from Precursor Cells 3 and *CA1^+^* colonocytes to *CEACAM7^+^*colonocytes were lost. Taken together, *CA2^+^* colonocytes consistently acted as the major CEACAM signaling initiators across all three donors, while *CEACAM7^+^* colonocytes were both initiators and receivers in Donors Y and Z. Notably, Precursor Cells 3 universally lost their signaling capacity after infection. Despite these commonalities, each donor exhibited distinct communication patterns within the CEACAM pathway, highlighting donor-specific differences in epithelial cell adhesion communications in response to infection-induced stress.

**Figure 5.**
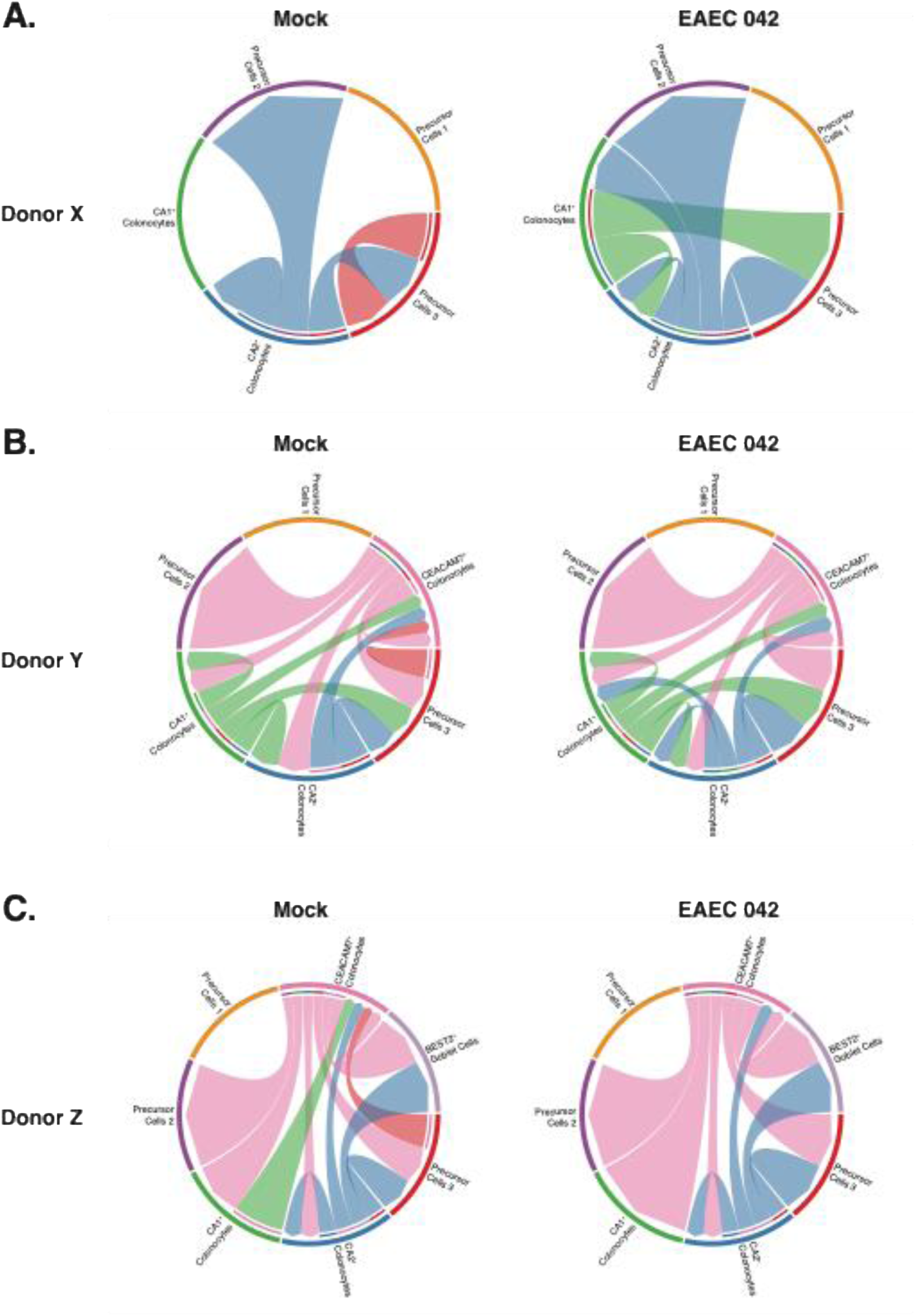
CellChat CEACAM Pathway Analysis. Chord diagrams showing the CEACAM pathway ligand-receptor interaction among colonoid cell clusters in Donor X **(A)**, Donor Y **(B)**, and Donor Z **(C)** under mock- or EAEC-infected conditions.

We next examined the growth and differentiation factor (GDF) pathway, in which GDF15 acts as the ligand and transforming growth factor beta receptor type 2 (TGFBR2) as the receptor. GDF15 is an inflammation-induced hormone, elevated in bacterial infections such as sepsis, where it has protective effects through enhancing tissue tolerance to inflammatory damage^26^. Ligand activation of TGFBR2 triggers canonical SMAD-mediated TGF-β signaling^27^, which reduces bacterial burden and mitigates tissue damage^28^. Thus, donor-specific GDF ligand-receptor interactions may provide insight into variability in bacteria-induced host inflammatory responses.

In Donor X (**Figure 6A**), GDF signaling occurred only within Precursor Cells 3 under mock conditions, but intensified after infection, with new interactions directed toward *CA2^+^* colonocytes. In Donor Y (**Figure 6B**), Precursor Cells 3 also dominated signaling under both mock and infected conditions, targeting their own cluster as well as all mature colonocyte populations. Infection further increased signaling, introducing new interactions such as communication from Precursor Cells 2 to *CEACAM7^+^* colonocytes and Precursor Cells 3. In Donor Z (**Figure 6C**), signaling was primarily initiated by Precursor Cells 3 and *BEST2*^+^ goblet cells, targeting most populations except Precursor Cells 1 and 2. Unlike the other donors, Donor Z showed a pronounced loss of signaling after infection. Notably, *CA1^+^* and *CEACAM7^+^* Colonocytes no longer act as receptors of interaction. Overall, Precursor Cells 3 consistently served as the major GDF ligand source across donors, highlighting its important role in mediating host inflammatory response. However, both the recipient populations and the infection-induced changes varied considerably, underscoring donor-specific differences in epithelial responses to bacterial infection.

**Figure 6.**
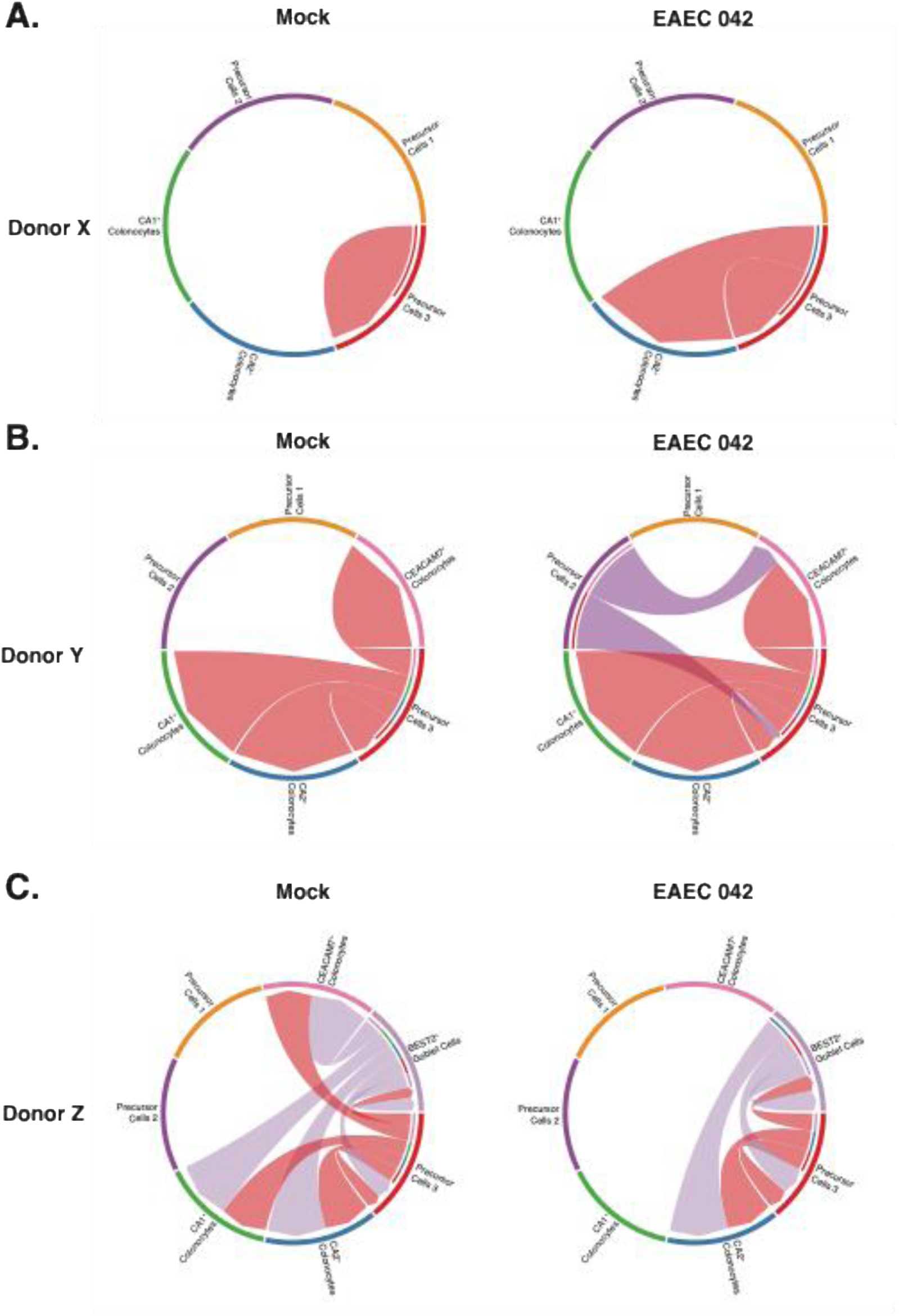
CellChat GDF Pathway Analysis. Chord diagrams showing the CEACAM pathway ligand-receptor interaction among colonoid cell clusters in Donor X **(A)**, Donor Y **(B)**, and Donor Z **(C)** under mock- or EAEC-infected conditions.

## DISCUSSION

The present study leverages the advanced technology of single-cell transcriptomics and the emerging tool of human organoid model systems to demonstrate the dramatic host variability in intestinal epithelial responses to EAEC infection. Building upon previous observations that EAEC infection displays a segment- and donor-dependent trend^15–16^, our study shows that such heterogeneity extends beyond bacterial adherence to cell type composition, transcriptional response, and intercellular communication. It emphasizes the urgent need to consider host genetic diversity when studying infectious disease mechanisms and developing therapeutic interventions.

Our single cell RNA-seq analysis shows that human colonoids from three healthy donors contain seven epithelial subpopulations, including three precursor clusters, three mature colonocyte clusters, and one goblet cell cluster (**Figure 2**). The first finding of this study is the remarkable baseline differences in colon epithelial cell composition among the three donors despite identical culturing conditions. Donor X was dominated by precursor populations, while Donors Y and Z were each enriched with mature colonocytes that have distinct gene signatures. These observations suggest that tissue stem cell-derived colon organoids may retain donor-specific characteristics that reflect the genetic landscape of the original tissue.

The striking heterogeneity in baseline cellular composition observed across donors likely underlies differences in infection susceptibility. Supporting this hypothesis, each donor exhibited distinct patterns of cellular remodeling following EAEC infection. The differential cellular responses to EAEC infection across donors reveal three distinct host phenotypes: hyperresponsive (Donor Y), intermediate responsive (Donor X), and non-responsive (Donor Z). This classification is supported not only by changes in cell type distribution (**Figure 2**) but also by the magnitude of transcriptional responses (**Figure 3**). For instance, after EAEC infection, Donor Y had a significant enrichment of *CA1^+^* Colonocytes and a robust upregulation of immune, damage response, and cellular growth pathways, which suggests an active epithelial defensive response. In contrast, despite similar bacterial load (**Figure 1**), Donor Z not only maintained a stable cellular composition, but also had minimal transcriptional changes after infection. It is uncertain if this relates to the selective upregulation of *CA2*^+^ Colonocytes as a result of the higher catalytic activity of CA2 in pH regulation. Mucin secretion by *BEST2*^+^ Goblet Cells in Donor Z might also help maintain homeostasis and protect the host from epithelial damage. The intermediate phenotype observed in Donor X, characterized by expansion of immune-activated Precursor Cells 3, may represent a balanced response that mounts antibacterial defense while minimizing tissue inflammatory damage. Furthermore, the donor-specific cell type distribution and remodeling further contribute to the highly heterogeneous DEG and pathway regulation in response to infection (**Figure 3**). The infection-induced DEGs of each donor not only differed in regulatory directions but also in the driving cell types: Donor X showed predominantly upregulated DEGs driven by Precursor Cells 2, Donor Y exhibited mixed regulation dominated by *CA1*⁺ Colonocytes, and Donor Z demonstrated predominantly downregulated DEGs driven by *CEACAM7*⁺ Colonocytes. Further analysis of overlapping DEGs across cell clusters revealed some conserved gene regulation signatures across colonocyte populations but mostly unique transcriptional responses. This suggests that epithelial subpopulations have evolved specialized roles in pathogen defense, with the distinct cellular architecture of each donor contributing to personalized antimicrobial response strategies. The GSEA pathway analysis provides mechanistic insights into the transcriptional changes induced by the pathogen. Interestingly, there was a universal upregulation of hypoxia pathways across multiple cell types in all three donors, reflecting universal metabolic stress induced by EAEC virulence factors. Nevertheless, the overall trend for pathway regulation is still unique to each donor. The heterogeneity in donor response to EAEC infection may explain why some individuals develop severe and chronic diarrhea while others remain asymptomatic even with similar bacterial exposure and colonization.

The CellChat analysis (**Figure 4**) further substantiated the donor-specific nature of EAEC epithelial responses by investigating the highly complex cell-cell communication networks. Given the high numbers of total upregulated and downregulated signaling events, it is remarkable that only a limited number of interactions were shared among the three donors. To our surprise, Precursor Cells 3 consistently acted as a central signaling cluster regardless of donors. It can be potentially explained by the universally high abundance of this cluster. Its expression of immune-activating genes might also contribute to its central role in coordinating tissue response to infection. However, in general, cell-cell communication networks varied considerably in complexity among donors. Notably, Donor Z exhibited the most intricate signaling network despite being a non-responder in transcriptional responses, suggesting that intercellular communication may be critical for maintaining epithelial homeostasis during infection. In addition, colon epithelial subpopulations exhibited donor-specific signaling patterns. For example, *CA2*⁺ Colonocytes downregulated communication with their own population and Precursor Cells 3 in Donor X, while these identical interactions were upregulated in Donors Y and Z. In-depth analysis of cell communication networks in the CEACAM (**Figure 5**) and GDF (**Figure 6**) pathways provided additional evidence on cell signaling events that are donor dependent. Consistent with cellular composition patterns, this heterogeneity is evident at both baseline and post-infection changes. CEACAM molecules are critical regulators of epithelial cell adhesion and have been implicated in inflammatory bowel disease pathogenesis. The donor-specific patterns of CEACAM signaling observed in our study may reflect variations in CEACAM expression in different colon epithelial clusters that can influence barrier function and lead to differential symptoms after pathogen exposure. The GDF15-TGFBR2 signaling axis is particularly relevant to bacterial infection responses, as GDF15 is known to promote tissue tolerance to inflammatory damage. The donor-specific differences in GDF15 signaling patterns suggest that the magnitude of inflammatory response to enteric infections may be dependent on donor cell type distributions and receptor expression profiles.

This study has some important clinical implications. First, the profound donor-specific differences we observed are important for considering precision medicine approaches when treating bacterial infections. The identification of distinct host response phenotypes highlights the importance of tailoring therapeutic interventions to individual needs. For example, hyperresponsive patients might require anti-inflammatory treatments, while non-responsive patients might benefit from a boost in antimicrobial response. Additionally, the vastly different host response despite comparable bacterial adherence suggests that current diagnostic approaches based on bacterial burden may not accurately predict disease severity. Instead, transcriptional signatures might serve as better prognostic indicators.

Our study has several limitations that should be considered when interpreting the findings. First, this study is limited to only three donor lines that varied in their age, sex, and ethnicity, which limits the generalizability of the identified response phenotypes. To verify these results, a large study that include donors with similar characteristics and genetic backgrounds should be conducted. Second, the HCO culture system is oversimplified due to the lack of innate and adaptive immune cells, which play crucial roles in pathogen clearance. Co-culture experiments that incorporate immune components will help capture a more comprehensive profile of host pathogen response. Nevertheless, considering the striking differences we observed across donors in this simplified system, we are certain that when combined with additional immune cells, this model will show additional complexity in donor-specific responses. Third, the 90-minute infection time only provides a snapshot of the acute phase of EAEC infection. Since this pathogen establishes long-term infections in children, time-course studies will be crucial to understand how donor-specific responses evolve over time and whether they predict long-term disease outcomes. However, the baseline differences that we observed are still striking regardless of the length of pathogen exposure. Lastly, since our study is limited to transcriptional level responses, it is difficult to verify that the heterogeneity in cell composition, gene regulation, and cell-cell communication is present at the protein level. Future investigations should integrate complementary approaches including single-cell proteomics and immunofluorescence imaging of cell subtype markers and key signaling molecules. These studies will be crucial for confirming whether donor-specific transcriptional signatures correspond to functional protein differences that impact infection outcomes.

## ACKNOWLEDGMENTS

This work was supported by NIH/NIAID under Grants CARB U19 AI157981-03, BCRC U19 AI116497-08, and GCID U19 AI144297-05. Tajhal D. Patel and Cristian Coarfa were partially supported by CPRIT RP210227 and RP200504, NIH/NCI P30 shared resource grant CA125123, NIH/NIEHS P42 ES027725 and P30 ES030285. The compute cluster used for analysis was partially supported by NIH 1S10OD032185.

## DATA AVAILABILITY

The single cell sequencing data will be electronically deposited and made publicly available.

## MATERIALS AND METHODS

### Human Colon Organoid Culture

Human colon organoids (HCOs) from healthy adult donors were cultured as previously described^15^. Briefly, HCO monolayers were generated from 3D enteroid cultures derived from healthy colon biopsy specimens. 3D colonoids were maintained in Matrigel and subsequently dissociated into single cells for seeding onto Matrigel-coated chambered slides (Greiner Bio-One, 543979). Cells were cultured in complete medium with growth factors and the ROCK inhibitor Y-27632 (Sigman, Y-0503) for initial attachment. After 24 h, the medium was replaced with differentiation medium, and cells were differentiated for 4 days with media changes every other day before infection.

### Bacteria Culture and Infection

EAEC prototype strain 042 (serotype 44:H18) was cultured overnight in LB broth at 37°C with aeration, then sub-cultured 1:20 in HCO differentiation media for 2 hours to obtain log-phase bacteria prior to infection. Day 4 differentiated HCO monolayers were then infected with EAEC 042 diluted in differentiation media at MOI=10 for 90 minutes at 37°C in a humidified incubator. Mock-infected HCO monolayers were treated with differentiation media without bacteria instead.

### Bacteria Enumeration

Infected HCO monolayers were washed with PBS and then dissociated using 100 µL/well of 0.05% Trypsin-EDTA (Gibco, 25300-054) at 37°C for 5 min. Cells were thoroughly dispersed by pipetting, diluted in PBS, and drip-plated onto LB agar. Plates were incubated overnight at 37°C, and colony-forming units (CFU) were counted the following morning.

### HCO Monolayer Collection for Single Cell RNA Sequencing

Mock- or EAEC-infected HCO monolayers on chamber slides were washed with PBS containing 5 mM EDTA and dissociated with 200 µL/well Accutase (STEMCELL Technologies, 07920). Cells were mixed using wide-bore pipette tips and incubated at 37°C for ∼55 min until fully digested. Wells from the same HCO donor line and treatment were pooled, neutralized with fetal bovine serum, and diluted with 1mL cold differentiation medium supplemented with ROCK inhibitor Y-27632 (Sigma, Y-0503). Then the cell suspensions were centrifuged at 300 × g for 5 min at 4°C, resuspended in 100 µL differentiation medium with Y-27632, and passed through Flowmi Cell Strainers (Millipore Sigma, BAH136800040). The resulting single-cell suspension was adjusted to 1,000 cells/µL for sequencing.

### Single Cell RNA Sequencing

Single-cell gene expression libraries were generated at the BCM ATC Single Cell Genomics Core using the Chromium Single Cell 3′ v3.1 kit (10x Genomics). Briefly, HCO single cell suspensions were combined with reverse transcription reagents, Gel Beads containing barcoded oligonucleotides, and oil on a Chromium X instrument (10x Genomics) to form Gel Bead-In-Emulsions (GEMs), enabling barcoded cDNA synthesis from each individual cell. Subsequently, the GEMs were disrupted and cDNA from each single cell was pooled. Following cleanup using Dynabeads MyOne Silane Beads, cDNA was amplified by PCR. The amplified cDNA was then fragmented, end-repaired, A-tailed, and ligated with sequencing adaptors. Final sequencing libraries were generated by amplification.

### scRNAseq Processing and Analysis

The 10x Genomics Cellranger software (v6.0) was used to demultiplex cell barcodes and map sequencing reads to the human reference genome (GRCh38). The resulting UMI counts matrix was further normalized using Scanpy (v1.8.1) where cells with UMI<500 were removed. Doublets were detected using the Scrublet (v0.2.3) package with an additional filtering removal of single cells containing greater than 8000 genes detected. Since the samples were derived from different patients, batch correction was performed using BBKNN (v1.6.0) to account for technical variation. The dimensionality of the dataset was reduced using principal component analysis, cells were clustered based on the leiden method and visualized by UMAP plots using Scanpy. Cell type annotation was accomplished using a combination of FindAllMarkers and FindTransferAnchors functions in Seurat (v5.0.3) along with published single cell annotation (PMID: 35176508, PMID: 34497389). Differentially expressed genes between infected and uninfected cells within each cluster and within each donor were identified using the FindMarkers function along with the Mann-Whitney Wilcoxon test where the significant threshold was FDR<0.1 and absolute fold change >1.25. GSEA was performed using the Hallmark collection and significance threshold of FDR<0.25. To identify developmental stages pseudotime trajectory analysis was performed using the slingshot R package (v.2.10.0) without setting a starting cluster (PMID: 29914354). Inference of cell-cell communication was determined using the CellChat R package (v.1.6.1) and the CellChatDB of human ligand-receptor interactions. Cell lineages and pseudotime were inferred using the Slingshot R package (v.2.10.0).

## SUPPLEMENTAL MATERIAL

**Supplementary Table 1.** Donor and Sample Information.

| Donor | Age | Sex | Race/Ethnicity | Colon Segment | Infection | Cell Count (Total) | Cell Count (Post Filtering) |
| --- | --- | --- | --- | --- | --- | --- | --- |
| X | 74 | Female | Caucasian | Ascending Colon | Mock | 2128 | 1286 |
|  |  |  |  |  | EAEC 042 | 3463 | 2300 |
| Y | 24 | Female | African American | Unknown | Mock | 10474 | 6258 |
|  |  |  |  |  | EAEC 042 | 13503 | 10463 |
| Z | 40 | Male | Caucasian | Transverse Colon | Mock | 11758 | 8874 |
|  |  |  |  |  | EAEC 042 | 13250 | 10170 |

**Supplementary Figure 1.**
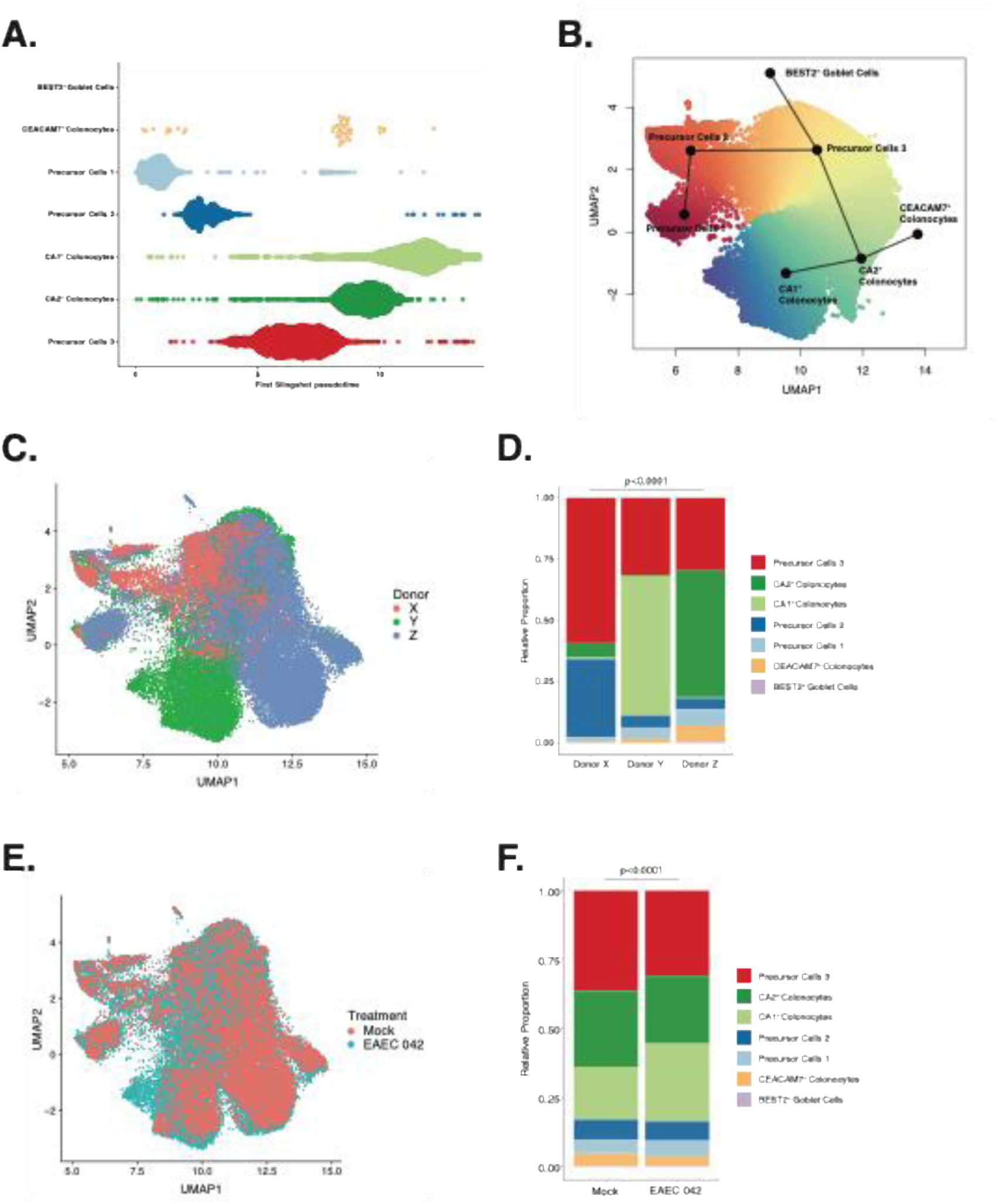
Cell type clustering and trajectory analysis. **(A)** Violin plot showing Slingshot pseudotime for each cell cluster. **(B)** UMAP showing Slingshot pseudotime trajectory analysis. **(C)** UMAP showing all cells separated by donors. **(D)** Bar plot depicting relative proportions of each cell cluster for each separate donor. **(E)** UMAP showing all cells separated by infection condition. **(F)** Bar plot depicting relative proportions of each cell cluster for mock- or EAEC-infected conditions.

**Supplementary Figure 2.**
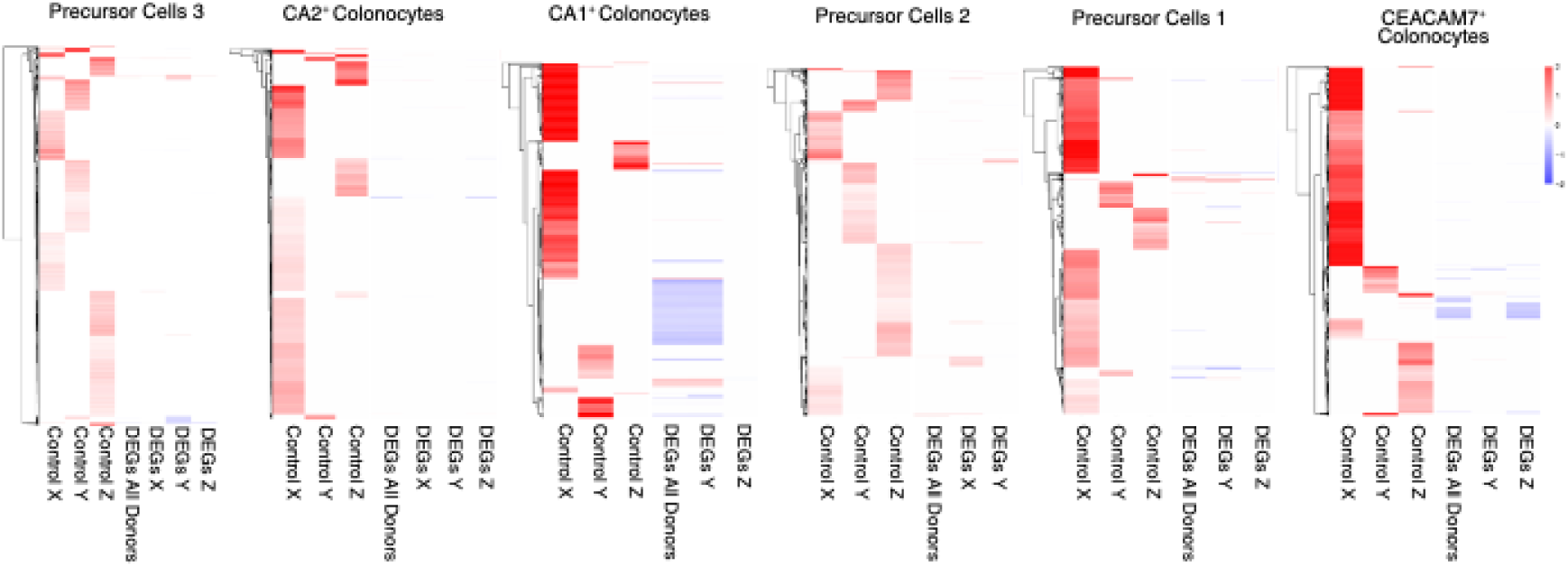
Heatmaps showing markers of mock-infected controls and differentially expressed genes (DEGs) induced by EAEC 042 infection.

## REFERENCES

1. Dehbanipour, R., and Ghalavand, Z. (2022). Anti-virulence therapeutic strategies against bacterial infections: recent advances. Germs 12, 262–275. 10.18683/germs.2022.1328.

2. Casadevall, A., and Pirofski, L. (2018). What Is a Host? Attributes of Individual Susceptibility. Infect Immun 86, e00636–17. 10.1128/IAI.00636-17.

3. Dutta, S., Pal, S., Chakrabarti, S., Dutta, P., and Manna, B. (1999). Use of PCR to identify enteroaggregative Escherichia coli as an important cause of acute diarrhoea among children living in Calcutta, India. J Med Microbiol 48, 1011–1016. 10.1099/00222615-48-11-1011.

4. Glandt, M., Adachi, J.A., Mathewson, J.J., Jiang, Z.D., DiCesare, D., Ashley, D., Ericsson, C.D., and DuPont, H.L. (1999). Enteroaggregative Escherichia coli as a cause of traveler’s diarrhea: clinical response to ciprofloxacin. Clin Infect Dis 29, 335–338. 10.1086/520211.

5. Wanke, C.A., Mayer, H., Weber, R., Zbinden, R., Watson, D.A., and Acheson, D. (1998). Enteroaggregative Escherichia coli as a potential cause of diarrheal disease in adults infected with human immunodeficiency virus. J Infect Dis 178, 185–190. 10.1086/515595.

6. Roche, J.K., Cabel, A., Sevilleja, J., Nataro, J., and Guerrant, R.L. (2010). Enteroaggregative E. coli (EAEC) Impairs Growth and Malnutrition Worsens EAEC Infection: A Novel Murine Model of the Infection Malnutrition Cycle. J Infect Dis 202, 506–514. 10.1086/654894.

7. Steiner, T.S., Lima, A.A., Nataro, J.P., and Guerrant, R.L. (1998). Enteroaggregative Escherichia coli produce intestinal inflammation and growth impairment and cause interleukin-8 release from intestinal epithelial cells. J Infect Dis 177, 88–96. 10.1086/513809.

8. Raju, B., and Ballal, M. (2009). Multidrug resistant enteroaggregative Escherichia coli diarrhoea in rural southern Indian population. Scand J Infect Dis 41, 105–108. 10.1080/00365540802641856.

9. Gonyar, L.A., Smith, R.M., Giron, J.A., Zachos, N.C., Ruiz-Perez, F., and Nataro, J.P. (2020). Aggregative Adherence Fimbriae II of Enteroaggregative Escherichia coli Are Required for Adherence and Barrier Disruption during Infection of Human Colonoids. Infect Immun 88, e00176–20. 10.1128/IAI.00176-20.

10. Paletta, A.C.C., Castro, V.S., and Conte-Junior, C.A. (2020). Shiga Toxin-Producing and Enteroaggregative Escherichia coli in Animal, Foods, and Humans: Pathogenicity Mechanisms, Detection Methods, and Epidemiology. Curr Microbiol 77, 612–620. 10.1007/s00284-019-018421.

11. Garcia, P.G., Silva, V.L., and Diniz, C.G. (2011). Occurrence and antimicrobial drug susceptibility patterns of commensal and diarrheagenic Escherichia coli in fecal microbiota from children with and without acute diarrhea. J Microbiol. 49, 46–52. 10.1007/s12275-011-0172-8.

12. Nüesch-Inderbinen, M.T., Hofer, E., Hächler, H., Beutin, L., and Stephan, R. (2013). Characteristics of enteroaggregative Escherichia coli isolated from healthy carriers and from patients with diarrhoea. Journal of Medical Microbiology 62, 1828–1834. 10.1099/jmm.0.065177-0.

13. Nataro, J.P., Deng, Y., Cookson, S., Cravioto, A., Savarino, S.J., Guers, L.D., Levine, M.M., and Tacket, C.O. (1995). Heterogeneity of enteroaggregative Escherichia coli virulence demonstrated in volunteers. J Infect Dis 171, 465–468. 10.1093/infdis/171.2.465.

14. Blutt, S.E., and Estes, M.K. (2022). Organoid Models for Infectious Disease. Annu Rev Med 73, 167–182. 10.1146/annurev-med-042320-023055.

15. Rajan, A., Vela, L., Zeng, X.-L., Yu, X., Shroyer, N., Blutt, S. E., Poole, N. M., Carlin, L. G., Nataro, J. P., Estes, M. K., Okhuysen, P. C., & Maresso, A. W. (2018). Novel Segment- and Host-Specific Patterns of Enteroaggregative Escherichia coli Adherence to Human Intestinal Enteroids. mBio, 9(1), 10.1128/mbio.02419-17. https://doi.org/10.1128/mbio.02419-17

16. Rajan, A., Robertson, M.J., Carter, H.E., Poole, N.M., Clark, J.R., Green, S.I., Criss, Z.K., Zhao, B., Karandikar, U., Xing, Y., et al. (2020). Enteroaggregative E. coli Adherence to Human Heparan Sulfate Proteoglycans Drives Segment and Host Specific Responses to Infection. PLOS Pathogens 16, e1008851. 10.1371/journal.ppat.1008851.

17. Liu, L., Saitz-Rojas, W., Smith, R., Gonyar, L., In, J.G., Kovbasnjuk, O., Zachos, N.C., Donowitz, M., Nataro, J.P., and Ruiz-Perez, F. (2020). Mucus layer modeling of human colonoids during infection with enteroaggragative E. coli. Sci Rep 10, 10533. 10.1038/s41598-020-67104-4.

18. Haber, A.L., Biton, M., Rogel, N., Herbst, R.H., Shekhar, K., Smillie, C., Burgin, G., Delorey, T.M., Howitt, M.R., Katz, Y., et al. (2017). A single-cell survey of the small intestinal epithelium. Nature 551, 333–339. 10.1038/nature24489.

19. Triana, S., Stanifer, M.L., Metz-Zumaran, C., Shahraz, M., Mukenhirn, M., Kee, C., Serger, C., Koschny, R., Ordoñez-Rueda, D., Paulsen, M., et al. (2021). Single-cell transcriptomics reveals immune response of intestinal cell types to viral infection. Molecular Systems Biology 17, e9833. 10.15252/msb.20209833.

20. Kuo, R.-L., Chen, Y.-T., Li, H.-A., Wu, C.-C., Chiang, H.-C., Lin, J.-Y., Huang, H.-I., Shih, S.-R., and Chin-Ming Tan, B. (2021). Molecular determinants and heterogeneity underlying host response to EV-A71 infection at single-cell resolution. RNA Biol 18, 796–808. 10.1080/15476286.2021.1872976.

21. Hebbelstrup Jensen, B., Olsen, K.E.P., Struve, C., Krogfelt, K.A., and Petersen, A.M. (2014). Epidemiology and Clinical Manifestations of Enteroaggregative Escherichia coli. Clin Microbiol Rev 27, 614–630. 10.1128/CMR.00112-13.

22. Traag, V.A., Waltman, L., and van Eck, N.J. (2019). From Louvain to Leiden: guaranteeing well-connected communities. Sci Rep 9, 5233. 10.1038/s41598-019-41695-z.

23. Jin, S., Guerrero-Juarez, C.F., Zhang, L., Chang, I., Ramos, R., Kuan, C.-H., Myung, P., Plikus, M.V., and Nie, Q. (2021). Inference and analysis of cell-cell communication using CellChat. Nat Commun 12, 1088. 10.1038/s41467-021-21246-9.

24. Saiz-Gonzalo, G., Hanrahan, N., Rossini, V., Singh, R., Ahern, M., Kelleher, M., Hill, S., O’Sullivan, R., Fanning, A., Walsh, P.T., et al. (2021). Regulation of CEACAM Family Members by IBD-Associated Triggers in Intestinal Epithelial Cells, Their Correlation to Inflammation and Relevance to IBD Pathogenesis. Front Immunol 12, 655960. 10.3389/fimmu.2021.655960.

25. Thomas, J., Klebanov, A., John, S., Miller, L.S., Vegesna, A., Amdur, R.L., Bhowmick, K., and Mishra, L. (2023). CEACAMS 1, 5, and 6 in disease and cancer: interactions with pathogens. Genes Cancer 14, 12–29. 10.18632/genesandcancer.230.

26. Luan, H.H., Wang, A., Hilliard, B.K., Carvalho, F., Rosen, C.E., Ahasic, A.M., Herzog, E.L., Kang, I., Pisani, M.A., Yu, S., et al. (2019). GDF15 is an Inflammation-Induced Central Mediator of Tissue Tolerance. Cell 178, 1231–1244.e11. 10.1016/j.cell.2019.07.033.

27. Lee, J., Fricke, F., Warnken, U., Schnölzer, M., Kopitz, J., and Gebert, J. (2015). Reconstitution of TGFBR2-Mediated Signaling Causes Upregulation of GDF-15 in HCT116 Colorectal Cancer Cells. PLOS ONE 10, e0131506. 10.1371/journal.pone.0131506.

28. Deng, Z., Fan, T., Xiao, C., Tian, H., Zheng, Y., Li, C., and He, J. (2024). TGF-β signaling in health, disease and therapeutics. Sig Transduct Target Ther 9, 61. 10.1038/s41392-024-01764-w.

